# Standardized Residue Numbering and Secondary Structure Nomenclature in the Class D β-Lactamases

**DOI:** 10.1101/2025.01.20.633977

**Authors:** Anastasiya Stasyuk, Clyde A. Smith

**Affiliations:** Stanford Synchrotron Radiation Lightsource, SLAC National Accelerator Laboratory, Menlo Park, CA; Department of Chemistry, Stanford University, Stanford, CA

**Keywords:** Class D β-lactamase, residue numbering, secondary structure annotation

## Abstract

Over 1370 class D β-lactamases are currently known, and they pose a serious threat to the effective treatment of many infectious diseases, particularly in some pathogenic bacteria where evolving carbapenemase activity has been reported. Detailed understanding of their molecular biology, enzymology and structural biology are critically important but the lack of a standardized residue numbering scheme and inconsistent secondary structure annotation has made comparative analyses sometimes difficult and cumbersome. Compounding this, in the post-AlphaFold world where we currently find ourselves, an extraordinary wealth of detailed structural information on these enzymes is literally at our fingertips, therefore it is vitally important that a standard numbering system is in place to facilitate the accurate and straightforward analysis of their structures. Here we present a residue numbering and secondary structure scheme for the class D enzymes and apply it to test targets to demonstrate ease at which it can be used.

---

The β-lactamases are bacterial enzymes which hydrolyze β-lactam antibiotics (penicillins, cephalosporins, monolactams, and carbapenems) and are a major contributor of acquired resistance to these therapeutic compounds. There are almost 8650 β-lactamase enzymes currently known^1^, divided into four molecular classes based on sequence similarity and conserved amino-acid motifs^2^. Classes A, C, and D use an active-site serine for catalysis, while class B enzymes are zinc-dependent metalloenzymes^3-5^. The serine β-lactamases all share two highly-conserved sequence motifs; SxxK at the N-terminus of an α-helix, containing the catalytic serine and a conserved lysine, and the K(S/T)G motif on a β-strand close to the active site. In the class D enzymes, the lysine in the SxxK motif is post-translationally modified by carboxylation and serves as a general base during catalysis. A third motif, SxV in a loop adjacent to the catalytic serine, is topologically common to all three classes (SDN in class A, YA(S)N in class C, and SxV in class D)^6^.

The earliest discovered class D enzymes (including OXA-1, OXA-2, OXA-5, and OXA-10) were from Gram-negative *Pseudomonas* and classified as narrow spectrum β-lactamases^7^. They were capable of hydrolyzing oxacillin more efficiently then benzylpenicillin^3^, hence the prevalent use of the OXA nomenclature. In the following years, OXA variants with a broader spectrum of activity (extended-spectrum β-lactamases) were identified^7^, and more alarmingly carbapenem-hydrolyzing enzymes, including OXA-23^8^ and OXA-24^9^ from *Acinetobacter baumannii*, and OXA-48 from *Klebsiella pneumoniae*^10^, have become widespread. A recent analysis of some of the earlier OXA enzymes has shown that even the narrow and extended spectrum enzymes produced resistance to carbapenems when expressed in *A. baumannii*^11^. Originally the class D β-lactamases were thought to be only present in the Gram-negative bacteria, but they have since been discovered also in Gram-positive bacteria (*Bacillus* and *Clostridioides*)^12-14^. There are currently over 1370 class D β-lactamases known^1^, and at least 46 variants have been structurally characterized. They all have the same overall three-dimensional architecture comprising two structural domains; a β-sheet domain with two flanking helices, and an all-α domain^15-16^, with the active site located between these two domains (Figure 1).

**Figure 1.**
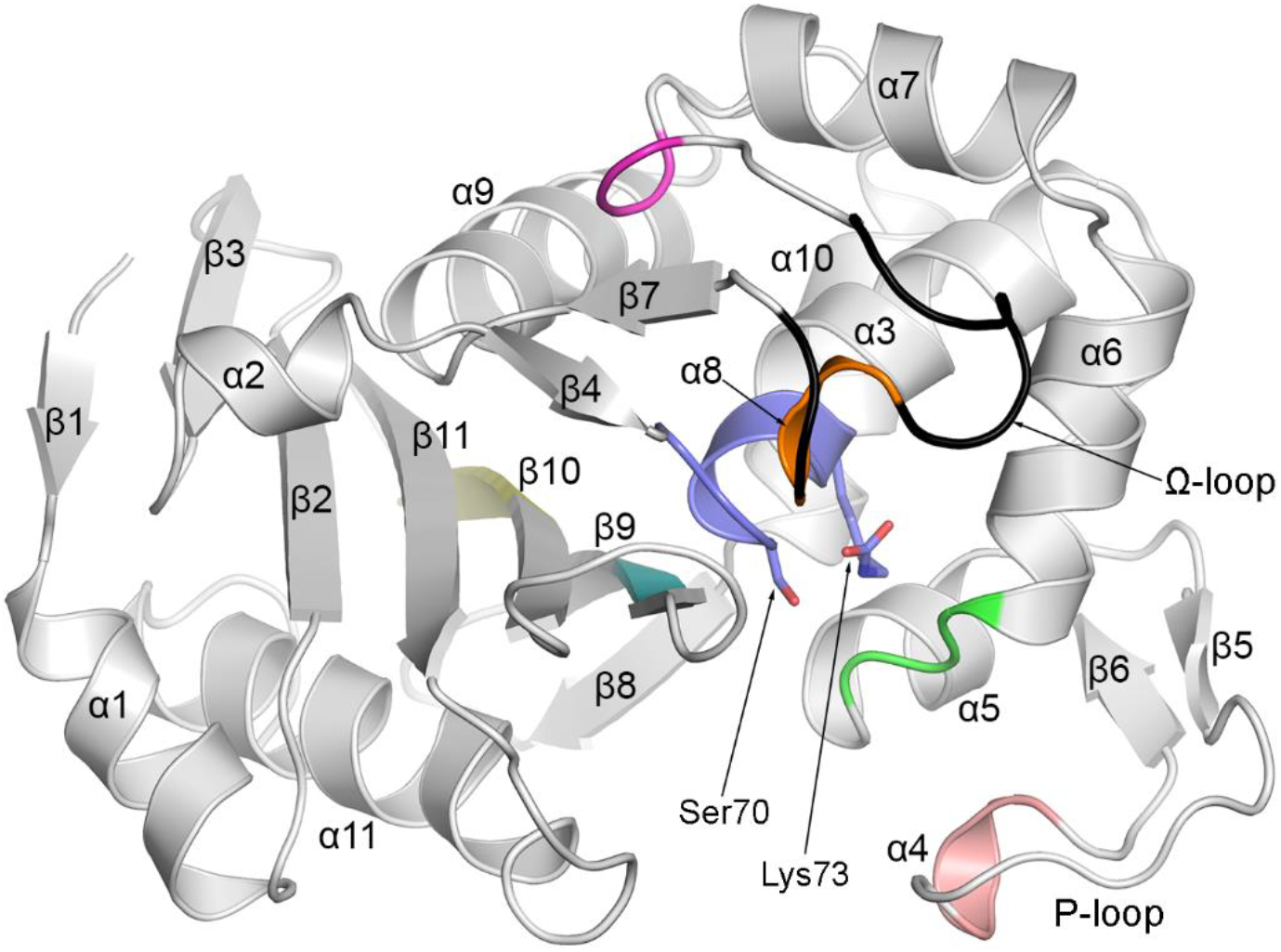
Ribbon diagram of OXA-48 as a representative class D β-lactamase structure. The DBL secondary structure annotation is shown. Two active site residues, Ser70 and carboxylated Lys73, are indicated. Sequence motifs specific to the class D enzymes are colored as follows: PASTFK (blue), FxxW (pink), SxV (green), YGN (magenta), FWL (orange), KTG (cyan) and GWxxGW (yellow). The Ω-loop is colored black.

Over the years standard amino acid numbering schemes for the class A and C serine β-lactamases^2, 17^ and the class B metallo-enzymes^18-20^ have been developed, and an early attempt was made to produce standard class D residue numbering based on comparison of five sequences with a number of class A enzymes^21^. Now that multiple structures of the class D enzymes from both Gram-negative and Gram-positive bacteria are known, time is past due to generate a robust numbering scheme that takes into account observed class D specific sequence motifs and sequence variations. We propose a numbering scheme (the DBL scheme to retain the name used earlier^21^) much like the Ambler scheme for the class A enzymes^2^, the BBL scheme for the class B enzymes^20^, and the SANC scheme for class C enzymes^17^, which can be used to refer to equivalent positions in the mature form of any class D β-lactamase. A standard secondary structure annotation scheme is also presented.

A multiple sequence alignment of all currently annotated class D β-lactamases from both Gram-negative and Gram-positive bacteria (Supporting Information File S1) confirmed the presence of the seven fingerprint motifs indicated in Figure 1. A second alignment of the Gram-negative OXA enzymes (Supporting Information File S2) was used to confirm familial relationships amongst the OXA enzymes (Table S1) and identify orphan OXAs with no family connections. An OXA family, defined as a group of two or more sequences with greater than 90% identity, is named according to the first enzyme identified. There are 21 OXA families comprising 11 or more members, another 29 families with 10 or fewer members, and 32 enzymes deemed orphans. The sequences of the parental members of each of the OXA families, along with the other (non-OXA) Gram-negative enzymes and the Gram-positive enzymes listed in the β-Lactamase DataBase (BLDB), were aligned and used to construct an unrooted phylogenetic tree (Figure S1). Not surprisingly the 19 *Acinetobacter* families clustered together in a single clade, as did the Gram-positive *Bacillus* enzymes, although the latter were quite separate from the *Clostridial* enzymes. Ultimately, to define the new DBL numbering and secondary structure scheme, a multiple alignment was generated using 25 unique sequences that include 12 OXA families, two orphans (OXA-45 and OXA-85), six other Gram-negative structures (not defined as OXAs), and five Gram-positive structures from the BLDB. These class D structures were superimposed on OXA-48 using the Secondary Structure Matching (SSM) algorithm^22^, and the sequence alignment adjusted manually based on equivalent amino acid positions (Figure S2).

To define the standardized secondary structure annotation for the new DBL scheme, reported annotations for the class D β-lactamases from 19 publications were collated (Table S2) and found to be somewhat haphazard. The nine major helices were generally consistently annotated, although two shorter 3_10_ helices were sometimes left out. The β-strand numbering was less consistent due to the intermittent identification of two regions where there are short interconnected antiparallel strands. To generate the DBL annotation, secondary structure assignments for the 25 structures were made using the STRIDE server^23^ and DSSP implemented in PROCHECK^24^ and WHAT-IF^25^. The assignments were then visually inspected and discrepancies from the algorithms were corrected in the annotations according to observed hydrogen bonding patterns and backbone torsion angles. This approach helped to identify short β-strands and β-bridges frequently missed (Figures 2A,B), and also corrected for the tendency of DSSP to under-predict strand and helix lengths^26^. The new DBL annotation is shown at the top of each sequence block in Figure S2. There are eleven helices (α1–α11) and eleven strands (β1–β11) (Table 1) joined by 14 short loops, and two longer loops which have more complex structures (Figure 1). The first is a long loop between helices α3 and α5 known as the P-loop^14, 27-28^, comprising two short anti-parallel strands (β5 and β6) linking the ends and a short 3_10_ helix (α4) (Figure 2C). The second is the Ω-loop between helices α7 and α9, which can have three different folds based upon the current analysis (Figure 2D). The most common type-I loop, observed in OXA-48 and several other enzymes (Figure S2), comprises a 3_10_ helix (α8) followed by a short strand (β7). In some enzymes there is what we define as a type-II loop with a six-residue insertion in the “OXA-48-like” type-I loop prior to α8. This type-II insertion juts outward from the Ω-loop, and helix α8 is longer than the equivalent helix in the shorter type-I loop (Figure 2D). The enzymes from *Clostridioides difficile* (CDD-1 and CDD-10) have what we define as a type-III loop, with a longer (∼10 residues) insertion in the “OXA-48-like” type-I sequence which folds more towards the active site and contains a second 3_10_ helix (α8’).

**Table 1.** DBL secondary structure, residue numbering and sequence motifs.

| Secondary structure<sup>a</sup> | Residue numbering<sup>a</sup> | DBL sequence motif |
| --- | --- | --- |
| strand $\beta 1^b$ | 25-28 | |
| helix $\alpha 1^c$ | 29-38 | |
| strand $\beta 2$ | 42-49 | |
| strand $\beta 3$ | 53-56 | |
| helix $\alpha 2$ | 59-63 | |
| strand $\beta 4$ | 65-67 | |
| helix $\alpha 3^d$ | 69-83 | PASTFK (68-73) |
| strand $\beta 5$ | 92-94 | |
| $3_{10}$ helix $\alpha 4$ | 103-105 | FxxW (103-106; P-loop) |
| strand $\beta 6$ | 108-110 | |
| helix $\alpha 5$ | 111-117 | |
| ($\alpha 5$-$\alpha 6$ loop) | | SxV (118-120) |
| helix $\alpha 6$ | 121-130 | |
| helix $\alpha 7$ | 132-142 | |
| ($\alpha 7$-$\alpha 8$ loop) | | YGN (144-146) |
| $3_{10}$ helix $\alpha 8$ | 156-158 | FWL (156-158; Ω-loop) |
| strand $\beta 7$ | 163-165 | |
| helix $\alpha 9$ | 166-178 | |
| helix $\alpha 10$ | 185-195 | |
| strand $\beta 8$ | 196-200 | |
| strand $\beta 9$ | 204-212 | KTG (208-210) |
| strand $\beta 10$ | 219-227 | GWxxGW (220-225) |
| strand $\beta 11$ | 232-240 | |
| helix $\alpha 11$ | 248-261 | |

<sup>a</sup> Residue numbering is based on the secondary structure elements in OXA-48. The length of these secondary structure elements may be different in other enzymes (refer to Figure S2).
<sup>b</sup> This strand is missing in some class D β-lactamase families, including the *Acinetobacter* OXAs.
<sup>c</sup> This helix is elongated in the *Acinetobacter* OXAs, and missing in the mature forms of other enzymes including OXA-45, OXA-57, OXA-427, AFD-1, ATD-1 and LoxA9.
<sup>d</sup> Helix $\alpha 3$ begins with a single turn of $3_{10}$ helix before starting the α-helix proper.

**Figure 2.**
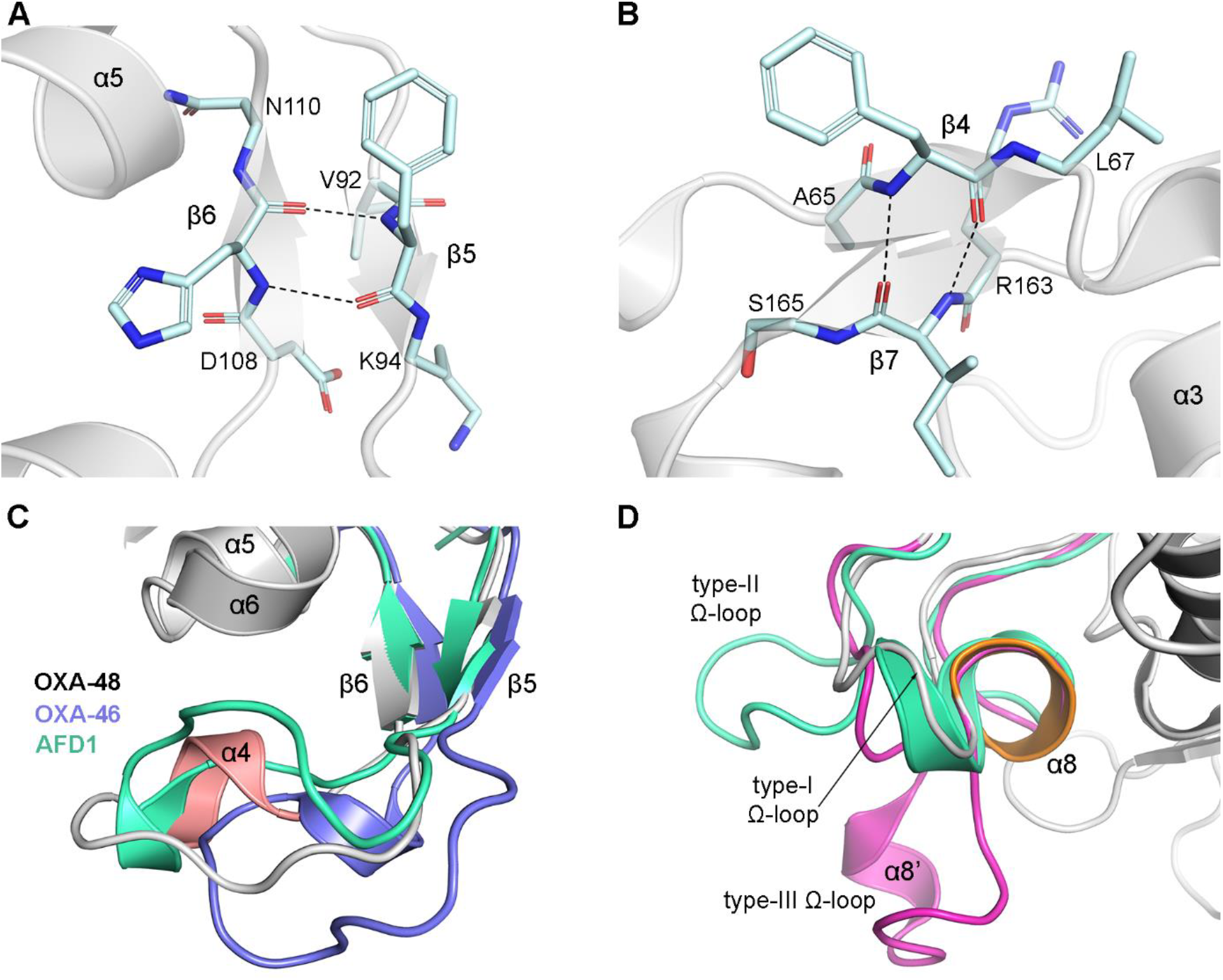
Some secondary structure elements in class D β-lactamases. **(A)** Antiparallel strands β5 and β6 at the beginning and end of the P-loop in OXA-48. **(B)** Antiparallel strands β4 and β7. Both sets of short strands in panels **A** and **B** are infrequently annotated in structure reports yet are important to the stability of their respective loops. Hydrogen bonds linking the strands are shown as black dashed lines. **(C)** Superposition of the P-loops of OXA-46 (blue) and AFD-1 (green) onto OXA-48 (gray). The helix α4 in OXA-48 is colored pink. **(D)** Superposition of the type-II Ω-loop of AFD-1 (green) and the type-III loop of CDD-1 (magenta) onto the more common short type-I loop of OXA-48 (gray). The helix α8 in OXA-48 is colored orange.

Standardization of the amino acid numbering in the new DBL scheme was based on OXA-48, as this enzyme has the most reported structures in the PDB. In the full OXA-48 sequence (the preprotein that includes the signal peptide) the catalytic serine is at position 70 with the carboxylated lysine at position 73, and the second universal serine β-lactamase motif (KTG) beginning at Lys208 (Figure S2). Utilizing the approach first used in the Ambler numbering scheme for the class A enzymes^2^ and subsequently adopted in the class B^20^ and class C^17^ schemes, in the aligned sequences, any residues missing relative to OXA-48 are skipped in numbering to maintain alignment with the OXA-48 numbering. Conversely, when there is an insertion in an aligned sequence relative to OXA-48, lower case letters are added to the residue number that just precedes the insertion. For example, the two longer type-II and type-III Ω-loops noted earlier are formed by insertions in two slightly different positions of the “OXA-48-like” type-I loop. The type-II insertion is between DBL positions 151 and 152 (numbered 151a through 151f, as in OXA-45 and others) (Figure S2). The type-III loop is inserted between DBL positions 153 and 154 and is numbered from 153a through 153j. Other insertions and deletions occur in loops between secondary structure elements and are consistent across the 25 structures (Table S3). The DBL numbering scheme is given at the bottom of each sequence block in Figure S2.

The alignment shown in Figure S2 is suitably robust to allow for its use in numbering and annotating any new class D structure. It is envisaged that any structures determined in the future will fall into one of four categories; (1) a member of an OXA family for which a structure is already known, (2) a member of an OXA family where there is no known structure, (3) orphan OXA enzymes and other Gram-negative class D β-lactamases (not defined as OXAs), or (4) novel Gram-positive enzymes. In the first category, it will be trivial to simply align the new sequence with that of the family member whose structure is known (to identify gaps or insertions) and assign the residue numbering and secondary structure for that family from Figure S2. Alternatively, the new structure could be superimposed with OXA-48, the sequences aligned, and any necessary adjustments made to give a true structure-based alignment. In the other three categories, the latter method would be adequate to give the correct DBL numbering and secondary structure annotation. To test the robustness of the proposed DBL scheme, we used AlphaFold2^29^ to predict the structures of some structurally uncharacterized class D enzymes, and applied the DBL scheme to these models. One example was taken from each category: (1) OXA-1042 from *Aeromonas veronii* (OXA-1 family), (2) OXA-213 from *Acinetobacter calcoaceticus* (a family with no structural representative), (3) OXA-919, an orphan from *Brucella pseudogrignonensis*; and (4) ClosD1, a Gram-positive enzyme from *Clostridium* sp. CH2. Signal peptides for the four enzymes were detected using the DeepSig server^30^ (Table S4) and removed prior to structure prediction. All structures were accurately predicted by AlphaFold2 as having a class D β-lactamase fold with high overall pLDDT and pTM scores (Figure S3 and Table S4). The predicted structures were superimposed on OXA-48 (Figures 3A-D) and individual pairwise sequence alignments between the target and OXA-48 were generated and manually adjusted based on the superpositions (Figures 4A-D). Secondary structure was determined with DSSP and STRIDE, followed by visual inspection (Figure S2).

**Figure 3.**
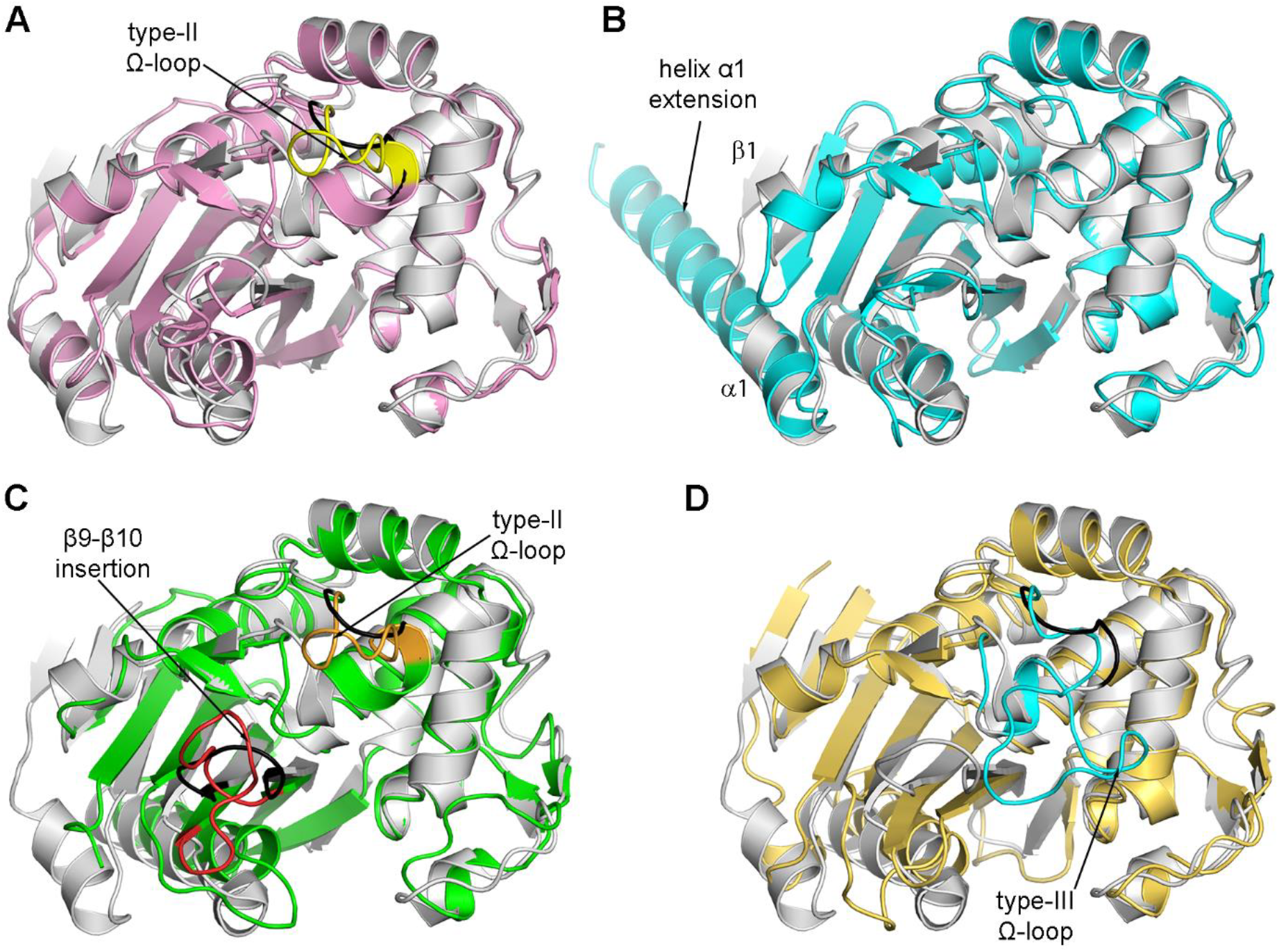
AlphaFold2 models of DBL test targets. The models are all superimposed on OXA-48 (gray ribbons). **(A)** OXA-1042 (pink), from the OXA-1 family. The type-II insertion in the Ω-loop (yellow) is indicated. The corresponding loop in OXA-48 is colored black. **(B)** OXA-213 (cyan), the parental member of the *Acinetobacter* OXA-213 family. The extended helix α1 in OXA-213 is indicated. **(C)** OXA-919 (green), an orphan enzyme. The type-II insertion in the Ω-loop (orange) and the 8-residue insertion in the β9-β10 loop (red) are indicated. The corresponding loops in OXA-48 are colored black. **(D)** ClosD1 (yellow) from *Clostridium* sp. CH2. The type-III extension in the Ω-loop (cyan) is indicated. The corresponding loop in OXA-48 is colored black.

**Figure 4.**
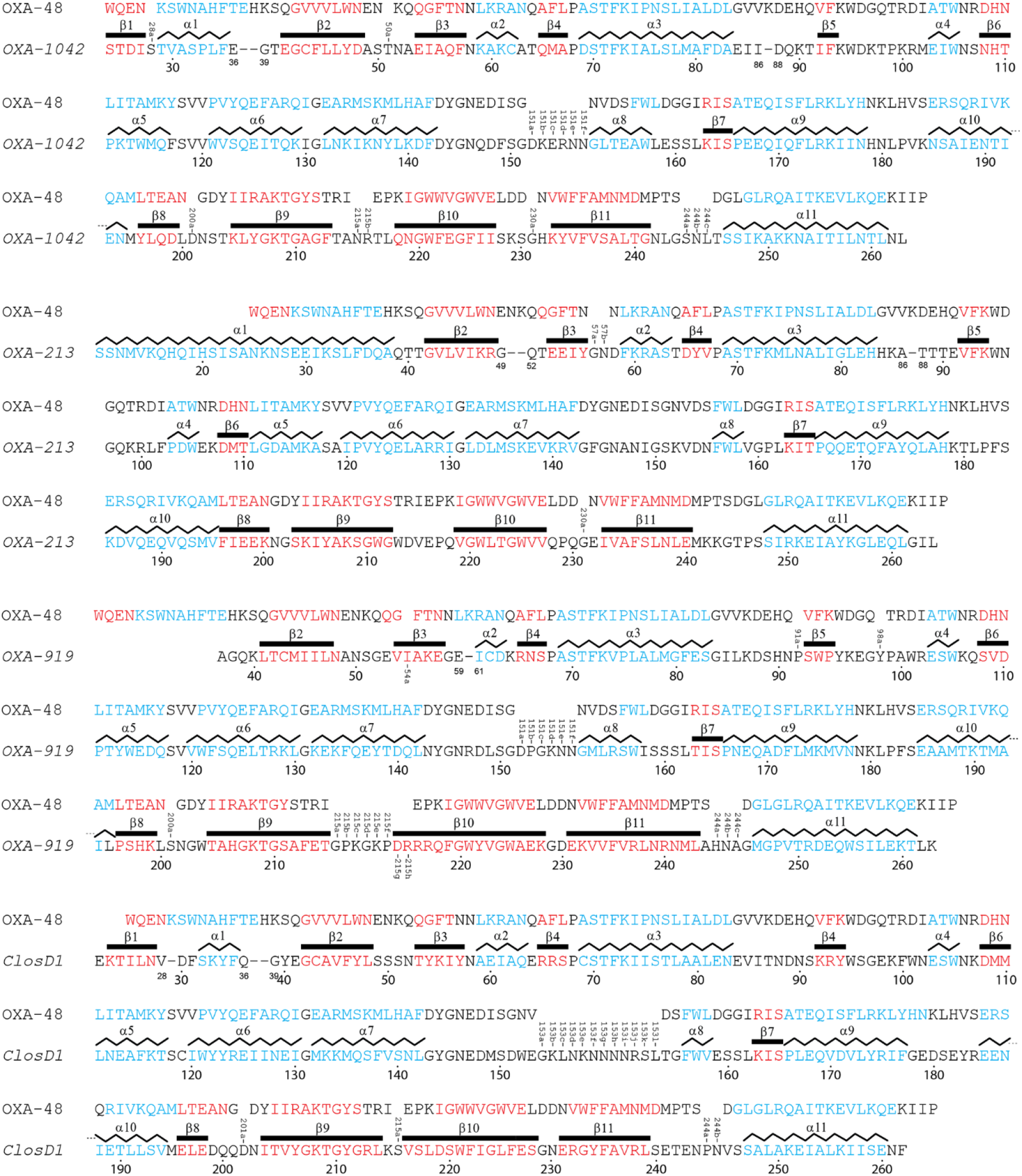
DBL residue numbering for the test targets. Structure-based sequence alignment of OXA-48 against **(A)** OXA-1042, **(B)** OXA-213, **(C)** OXA-919, and **(D)** ClosD1. In all alignments, the mature form of the target sequence is used. The DBL numbering scheme is given under each section. Secondary structure assignment is given for each predicted model. Amino acids in helices are colored cyan and those in strands are colored red. Insertions in the target sequences are indicated as gaps in the OXA-48 sequence with lower-case letters added to the numbers. Deletions in the target sequences are shown as dashes with the numbering on each side of the deletion. In OXA-213, β1 is missing and is skipped in the sequential annotation of the secondary structure elements. In OXA-919, strand β1 and helix β1 are missing and secondary structure annotation begins with strand β2.

The main difference between OXA-1042 (category 1) and OXA-48 is in the Ω-loop, where there is predicted to be a type-II insertion after DBL position 151 (Figures 3A), topologically equivalent to type-II loop extensions in OXA-1 and AFD-1 (Figure S4A). There are smaller 1-3 residue insertions at DBL positions 28, 50, 200, 215, 230 and 244, and two small deletions (37-38 and 87). As expected, the OXA-1042 model closely resembles the known structure of the OXA-1, and DBL numbering applied to OXA-1042 is identical to the parental enzyme (Figure 4A).

In the predicted OXA-213 model (category 2), helix α1 is five turns longer than in OXA-48 and there is no strand β1 (Figure 3B). The remainder of the structure is similar to OXA-48 with two minor insertions at positions 57 and 230, and two deletions (50-51 and 87) (Figure 4B). The elongated helix α1 is also seen in crystal structures of OXA-23, OXA-24, OXA-51, OXA-58 and OXA-143^31-35^, and superposition of the OXA-213 model onto OXA-23 shows that the predicted α1 extension is isostructural with helix α1 in the OXA-23 structure (Figure S4B). The similarities at the N-terminus are not unexpected since all enzymes are derived from *Acinetobacter* spp. and share ∼55-80% identity, and it is expected that a long helix α1 will be a feature of all *Acinetobacter* OXAs. Indeed, AlphaFold2 models of the parental sequences of the other twelve *Acinetobacter* families all show an extended N-terminal helix (Figure S5).

OXA-919 (category 3) is an orphan with only minimal similarity with other class D enzymes (19-43% sequence identity), and the model is predicted to be divergent from OXA-48 in several places. The mature form of OXA-919 lacks the first two secondary structure elements (β1 and α1) (Figure 4C), thus resembling several of the class D structures (Figure S2). There is a type-II insertion in the Ω-loop which is isostructural with the insertions in OXA-1 and AFD-1 (Figure S4A), in addition to small insertions at DBL positions 54, 91, 98, 200 and 244, and one deletion (60) (Figure 4C). Finally, there is an insertion of 8 residues in the β9-β10 loop, reminiscent of several enzymes (including OXA-57 and OXA-427) (Figure S2), and this region in the class D enzymes has been implicated in substrate recognition^14, 36-37^. The DBL numbering for this enzyme is shown in the alignment in Figure 4C, with secondary structure annotation beginning at strand β2 to coincide with the DBL scheme.

Finally, in the ClosD1 model (category 4), the major difference compared to OXA-48 is in the Ω-loop, which is nine residues longer in the *Clostridium* enzyme (Figures 3D). Although it bears some similarity to the type-III Ω-loop in the CDD-1 and CDD-10 crystal structures, there is no second 3_10_ helix predicted and the loop is essentially unstructured. In addition, there are small insertions at DBL positions 201, 215 and 244, and two deletions (29 and 37-38) (Figure 4D). In all four test targets, the DBL scheme outlined in Figure S2 was readily able to provide standardized numbering for these novel enzymes, with insertions and deletions generally limited to loops between secondary structure elements.

The implementation of standardized residue numbering and secondary structure schemes for the class D β-lactamases is long overdue and will prove beneficial for the quick and accurate comparison of their structures. In our own experience we have encountered situations where a single amino acid position was compared across several structures and the use of preprotein residue numbering for each structure made the descriptions very clumsy and unwieldy in both the text and the figures. In practical terms, it will be important that authors who employ the DBL scheme, explicitly state this early in their papers. We suggest that at first mention of an amino acid position, it should be given as both the preprotein number and the DBL equivalent, then the DBL scheme can be used to easily compare different enzymes and identify key residues involved in function and substrate binding. Secondary structure should be annotated as shown in Figure 1 and Figure S2, and any missing elements should be skipped so that subsequent elements coincide with the DBL scheme. The universal adoption of the standardized numbering schemes for the classes A, B, and C β-lactamases has greatly simplified their description, and it is our hope that acceptance of the DBL scheme for the class D enzymes will streamline subsequent structural and functional analyses of these enzymes.

## Methods

### Class D β-lactamase sequence alignments

Amino acid sequences for class D OXA β-lactamases were identified from the BLDB and downloaded from GENBANK (www.ncbi.nlm.nih.gov/genbank/). Additional OXA enzymes not listed in the BLDB were also found in GENBANK and downloaded. Multi-sequence alignments were performed using Clustal Omega^38^ (www.ebi.ac.uk/jdispatcher/msa/clustalo) on all the sequences, and on the OXA sequences separately. The results were downloaded as alignment files suitable for visualization with JALVIEW^39^. The sequences of the parental members of each of the 50 OXA families, eleven Gram-positive enzymes and 9 other Gram-negative enzymes were aligned with MAFFT (v.7)^40^ (mafft.cbrc.jp/alignment/server/) and the results used to generate a phylogenetic tree which was rendered using the iTOL server^41^ (itol.embl.de/).

### Structure predictions

AlphaFold2 structure predictions were run on the ColabFold server (v.1.5.5) (colab.research.google.com/github/sokrypton/ColabFold/blob/main/AlphaFold2.ipynb) using no structural template information. The top 5 ranked models were relaxed with Amber. The *Clostridium* sequence (BRC ID 2949990.6.peg.857) was found by searching the Bacterial and Viral Bioinformatics Resource Center (www.bv-brc.org). The enzyme was designated ClosD1 for this paper, and it shares 40% identity with CDD-1 and CDD-2.

All structural figures were generated with PyMOL (v2.5.2) from Schrodinger.

## Supporting information

Sequence alignment file 1

Sequence alignment file 2

## Supporting Information

Class D β-lactamase phylogenetic tree, structure-based sequence alignment of 25 class D β-lactamases, AlphaFold2 results, DBL test targets, AlphaFold2 models of 12 *Acinetobacter* OXA families, OXA enzyme families, secondary structure assignment for class D β-lactamases, DBL insertions and deletions, DBL test targets (PDF)

Supporting Information File 1, sequence alignment of all class D β-lactamases (txt)

Supporting Information File 2, sequence alignment of all OXA enzymes (txt)

## Acknowledgements

The SSRL Structural Molecular Biology Program is supported by the DOE Office of Biological and Environmental Research, and by the National Institutes of Health, National Institute of General Medical Sciences (P30GM133894). The contents of this publication are solely the responsibility of the authors and do not necessarily represent the official views of NIGMS or NIH. AS was supported by the Science Undergraduate Laboratory Internships (SULI) program, Department of Energy, Office of Science.

## Abbreviations

DBL: standardized class D β-lactamase residue numbering and secondary structure annotation scheme
BLDB: Beta-Lactamase DataBase

---

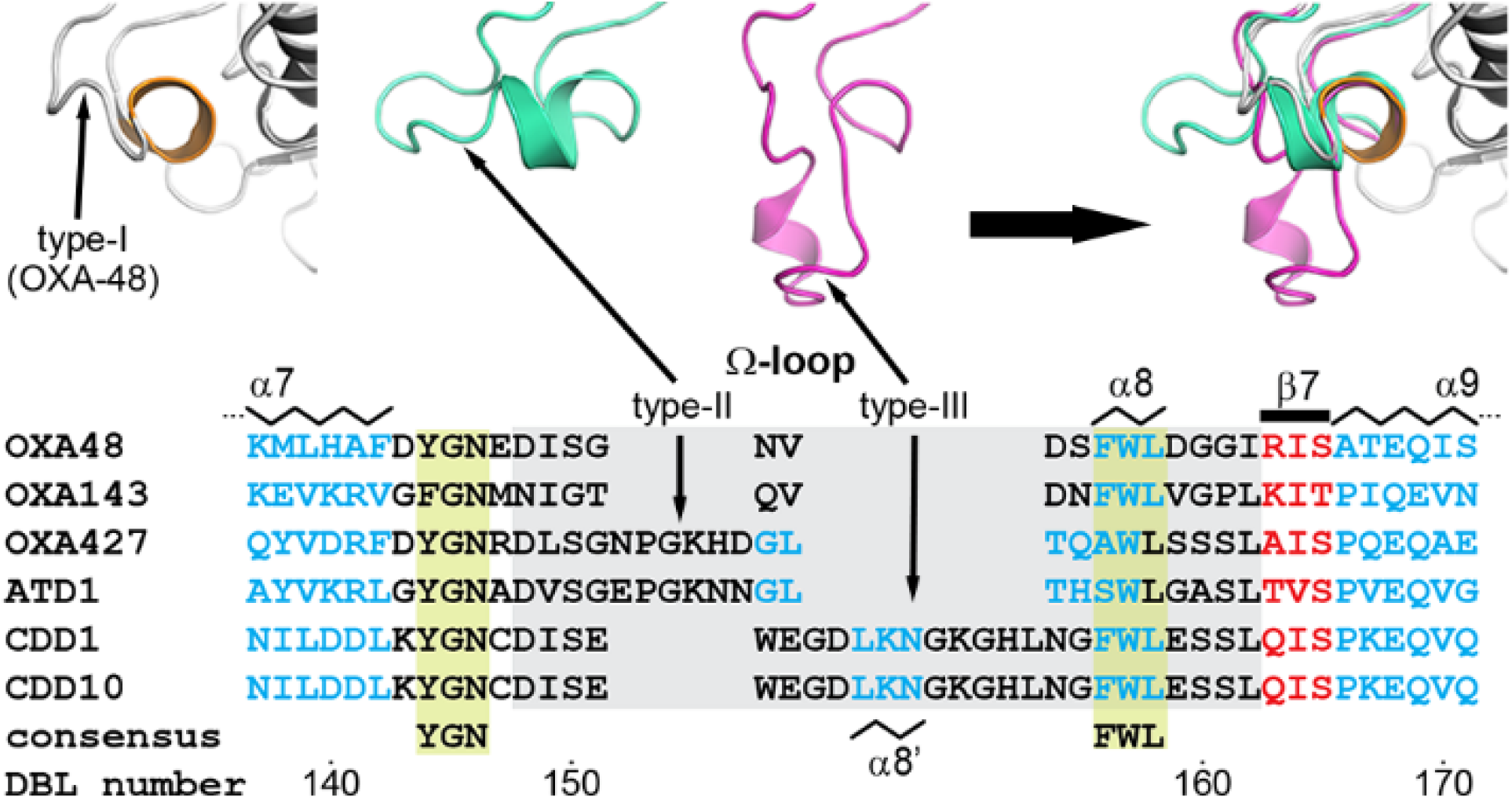

**TOC graphic**

**For Table of Contents only**

