## Supporting Information for "Standardized Residue Numbering and Secondary Structure Nomenclature in the Class D β-Lactamases"

### Supplemental Figures

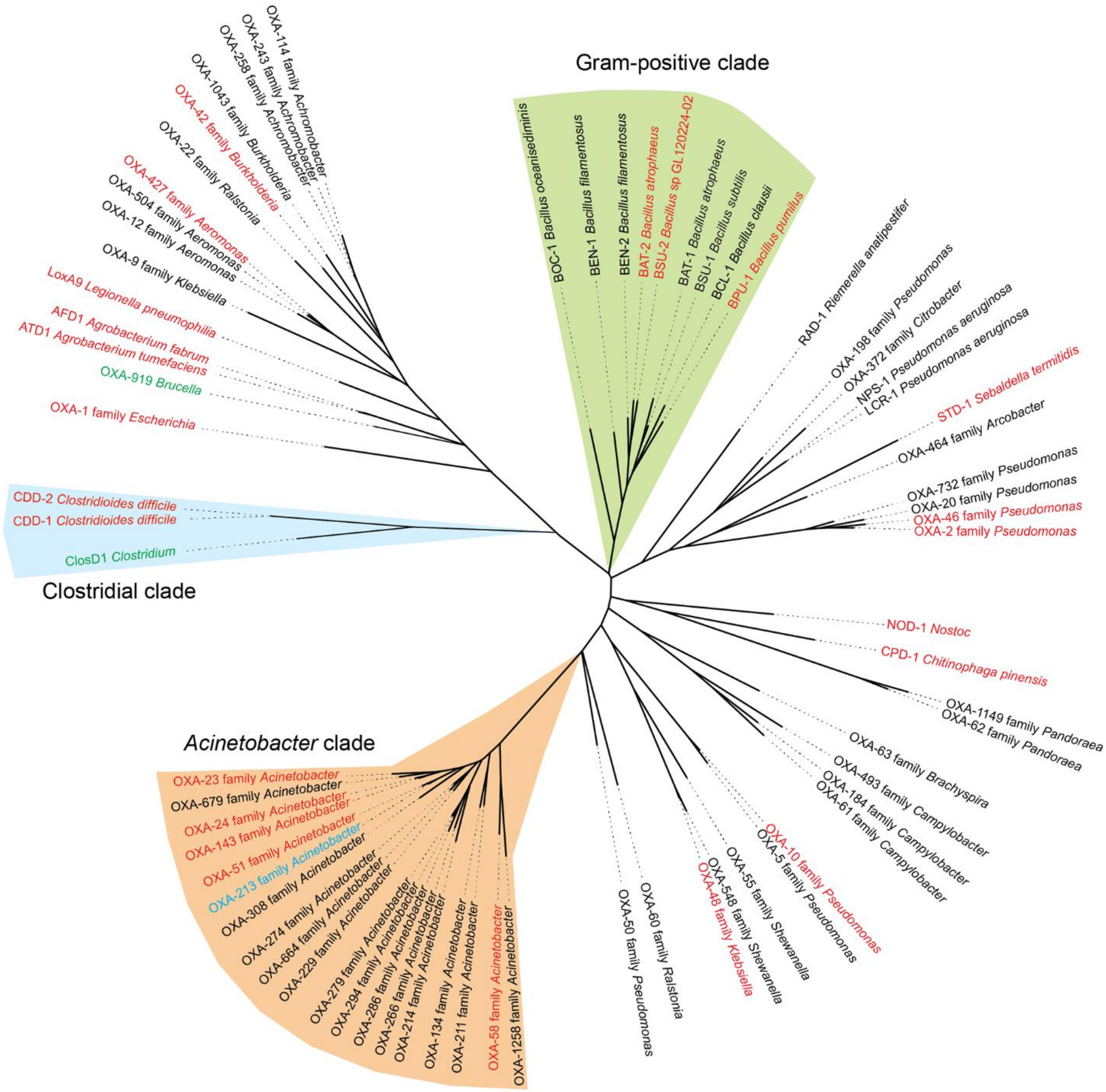


**Figure S1.** Phylogenetic tree from a multiple sequence alignment of the class D enzyme families. The Gram-positive *Bacillus* clade is shaded pale green, the *Acinetobacter* clade is shaded light orange, and the Clostridial clade is shaded light blue. Those families where X-ray structures of at least one member have been determined are colored red. The target sequences chosen to test the DBL scheme are colored green (OXA-213, OXA-919 and ClosD1). The fourth target (OXA-1042) is a member of the OXA-1 family.

```
                        β1      α1         β2            β3        α2     β4        α3
OXA48             KEWQEN KSWNAHFTEHKSQGVVVLWNEN  KQQG FTN  NLKRANQAFLPASTFKIPNSLIALDLGV
OXA1                STDISTVASPLFE--GTEGCFLLYDAS  TNAEIAQF  NKAKCATQMAPDSTFKIALSLMAFDAEI
OXA2             QEGTLER SDWRKFFSEFQAKGTIVVADERQADRAM LVF  DPVRSKKRYSPASTFKIPHTLFALDAGA
OXA10              SITEN TSWNKEFSAEAVNGVFVLCKSS  SKSC ATN  DLARASKEYLPASTFKIPNAIIGLETGV
OXA23    HNLINETPSQIVQGHNQVIHQYFDEKNTSGVLVIQTD-  -KKI NLYGNALSRANTEYVPASTFKMLNALIGLENQK
OXA24  SIKTKSEDNFHISSQQHEKAIKSYFDEAQTQGVIIIKEG-  -KNL STYGNALARANKEYVPASTFKMLNALIGLENHK
OXA45                              QMLECTLVADAA  SGQELYRK  GA--CDKAFAPMSTFKVPLAVMGYDAGI
OXA46            QEHVVIR SDWKKFFSDLQAEGAIVIADERQAKHTL SVF  DQERAAKRYSPASTFKIPHTLFALDADA
OXA51       NPNHSASKSDEKAEKIKNLFNEVHTTGVLVIQQG-  -QTQ QSYGNDLARASTEYVPASTFKMLNALIGLEHHK
OXA57                              AKTICTAIADAG  TGKLLVQD  GD--CGRRASPASTFKIAISLMGYDAGF
OXA58   SRAKTSTIPQVNNSIIDQNVQALFNEISADAVFVTYDG-  -QNI KKYGTHLDRAKTAYIPASTFKIANALIGLENHK
OXA85           IISFGNEN QFMKEIFERKGLNGTFVVYDLK  NDKI DYY  NLDRANERFYPASSFKIFNTLIGLENGI
OXA143  IQTKFEDTFHTSNQQHEKAIKSYFDEAQTQGVIIIKKG-  -KNI STYGNNLTRAHTEYVPASTFKMLNALIGLENHK
OXA427                              ATGCMLFADG-  SGKPFSAQ  GD--CASQLPPASTFKIPLALMGYDSGF
AFD1                           QQQPQAFECTLVTSIE  TGAIINQQ  GA--CDQRVAPASTFKVPLALIGFDAGI
ATD1                            QQSQAFECTLVTSIE  TGAVINQQ  GA--CDQRVAPASTFKVPLALIGYDAGI
CPD1           CAPNNVENE KSWEKYFAEYKVEGCFMLFNNS  QGTF KVY  NLERSQQRFLPASTFKIFNSLVGLETGV
LoxA9                               QDTCFLAKEN-  -QTVLKRE  GN-DCDQRYSPASTFKIALSLMGFDSGI
NOD1         QEIQTPAIPVN VNLGRSFNQLGIKGSILIYDRN  NKKF YEH  NAARNSQSFLPASTFKIFNSLVALETGV
STD1                   N VNLQKIFDENKITGSVTIYDYK  NKIW IYS  NEEDSKIRRLPASTFKIPNSLIFLEEEV
-------------------------------------------------------------------------------------
BAT2         KMPGEKSVNVSKLNVDEFFK--DRDGTFILHDVQ  KDKT FIY  NESRAKERQTPQSTFKVPNALIGLQVKA
BSU2        ASSAHEKHLNVSKMNVDDEFK--DTDGTFILHDLQ  KDQT FVY  NRKRANQRQTPQSTFKVVNALIGLQVKA
BPU1                   EARSIAWSVDEFFK--NREGTFVIQEVK  EKSP WVY  NKKRAKERFAPQSTFKVANALIGLQTGA
CDD1 NKSKIKNNDKNKKEAVNIV -DYSDCFE--GISGGAIFCNTK  NKEY NIY  NKELIETRRSPCSTFKIVSTLIGLEKGV
CDD10                 MV -DYSDCFE--GISGGAIFYNTK  NKEY NIY  NKELIETRRSPCSTFKIVSTLIGLEKGV
consensus                                                            PASTFK
                        -28a                 -50a -50b -54a -57a -57b
DBL number                30        40        50             60        70        80
```

Gram-negative (OXA48 – STD1); Gram-positive (BAT2 – CDD10)

```
                β5  P-loop   α4   β6    α5         α6          α7              Ω-loop
OXA48     VKDEHQ VFKWDGQTRD  IATWNRDHNLITAMKYSVVPVYQEFARQIGEARMSKMLHAFDYGNEDISG      NV
OXA1      I-DQKT IFKWDKTPKG  MEIWNSNHTPKTWMQFSVVWVSQEITQKIGLNKIKNYLKDFDYGNQDFSGDKERNNGL
OXA2      VRDEFQ IFRWDGVNRG  FAGHNQDQDLRSAMRNSTVWVYELFAKEIGDDKARRYLKKIDYGNADPST      SN
OXA10     IKNEHQ VFKWDGKPRA  MKQWERDLTLRGAIQVSAVPVFQQIAREVGEVRMQKYLKKFSYGNQNISG      GI
OXA23     T-DINE IFKWKGEKRS  FTAWEKDMTLGEAMKLSAVPVYQELARRIGLDLMQKEVKRIGFGNAEIGQ      QV
OXA24     A-TTNE IFKWDGKKRT  YPMWEKDMTLGEAMALSAVPVYQELARRTGLELMQKEVKRVNFGNTNIGT      QV
OXA45     LVDAHNPRWDYKPEFNG  YKFQQKTTDPTIWEKDSIVWYSQQLTRKMGQKRFAAYVAGFGYGNGDISGEPGKSNGL
OXA46     VRDEFQ VFRWDGVNRS  FAGHNQDQDLRSAMRNSTVWVYELFAKDIGEDKARRYLKQIDYGNVDPST      IK
OXA51     A-TTTE VFKWDGQKRL  FPEWEKDMTLGDAMKASAIPVYQDLARRIGLELMSKEVKRVGYGNADIGT      QV
OXA57     LRNEHDPVLPYRDSYIAWGGEAWKQPTDPTRWLKYSVVWYSQQVAHHLGAQRFAQYAKAFGYGNADVSGDPGQNNGL
OXA58     A-TSTE IFKWDGKPRF  FKAWDKDFTLGEAMQASTVPVYQELARRIGPSLMQSELQRIGYGNMQIGT      EI
OXA85     VKNVDEMFYYYDGSKVF  LDSWAKDSNLRYAIKVSQVPAYKKLARELGKERMQEGLNKLNYGNKEIGS      EI
OXA143    A-TTTE IFKWDGKKRS  YPMWEKDMTLGDAMALSAVPVYQELARRTGLDLMQKEVKRVGFGNMNIGT      QV
OXA427    LVDEQLPALPFKAGDPDF LPEWKQTTTPSRWMTYSVIWYSQRLTEWLGAARFQQYVDRFDYGNRDLSGNPGKHDGL
AFD1      LQDGKTPAWDWKPGTEA  RASDRKTVDPTIWEQDSVLWYSREITRRLGPEKFAAYVKRLGYGNADVSGEPGKNNGL
ATD1      LLDDKTPAWDWKPGTEA  RAQDRKTVDPTIWEQDSVLWYSRELTRRLGPEKFAAYVKRLGYGNADVSGEPGKNNGL
CPD1      IKDTSF VIPWDGVTRD  MPEWNHDLSMQQAFRVSAVPYFQEVARRITKPVMQHWLDTVKFGNMKI-S      KI
LoxA9     LKDELHPEWPYKKEYELY LNVWKYPQNPHTWIRDSCVWYSQALTRQLGMKRFKGYVDAFHYGNQDVSGDKGQNNGL
NOD1      ISNDVA ILTWDGMQRQ  FPTWNQDTNIRQAFRNSTVWFYQVLARKIGHERMEKFIKQVGYGNLQIGT    PEQI
STD1      VKDENE AMEWDGIKRY  IENWNKDLNLREAYEYSALWFYMKGAGKIKSEKYKEYLKEFNYGNQIVSE      KK
-------------------------------------------------------------------------------------
BAT2      VRDEYD VKRWDGTERE  FESWNRDHTLGSAMRDSVIWYYQAMARDIGEDRMKDWLHRISYGNEDISG      GI
BSU2      VRDEYD VKRWDGVKRE  FESWNRDHTLGSAMRESAIWYYQALARDIGEERMKTWLHTLSYGNEDISG      GI
BPU1      VRDEYD IKYWDGVKRE  IDNWNRDHTLGSGMRDSVVWYYQAMARDIGEERMNHWVKAIHYGNKDISG      GI
CDD1      INSKES VMGYDGTDYP  NKNWNKNLSLEEAFKESCVWYYKKLINKVDAKSVQNILDDLKYGNCDISE      WE
CDD10     INSKES VMGYDGTEYP  NKNWNKNLSLEEAFKESCVWYYKKLIDKVDAKSVQNILDDLKYGNCDISE      WE
consensus                    FxxW            SxV                       YGN
                -91a     -101a -101b                                        -151a -151b -151c -151d -151e -151f
DBL number    90         100         110       120       130       140       150 type-II
```

Gram-negative (OXA48 – STD1); Gram-positive (BAT2 – CDD10)

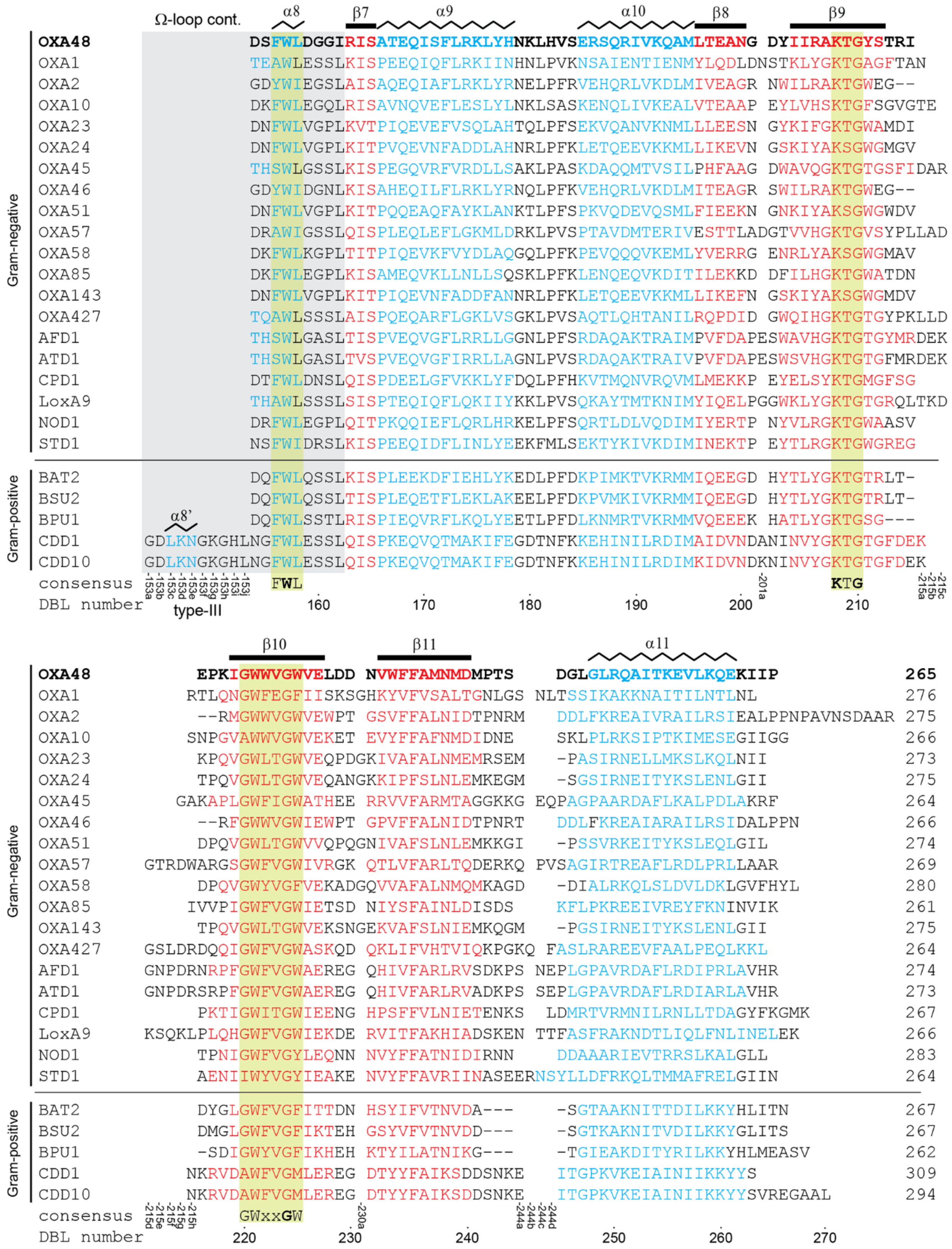

Ω-loop cont.
α8 β7 α9 α10 β8 β9
Gram-negative
OXA48 DSFWLDGGIRISATEQISFLRKLYHNKLHVSERSQRIVKQAMLTEANG DYIIRAKTGYSTRI
OXA1 TEAWLESSLKISPEEQIQFLRKIINHNLPVKNSAIENTIENMYLQDLDNSTKLYGKTGAGFTAN
OXA2 GDYWIEGSLAISAQEQIAFLRKLYRNELPFRVEHQRLVKDLMIVEAGR NWILRAKTGWEG--
OXA10 DKFWLEGQLRISAVNQVEFLESLYLNKLSASKENQLIVKEALVTEAAP EYLVHSKTGFSGVGTE
OXA23 DNFWLVGPLKVTPIQEVEFVSQLAHTQLPFSEKVQANVKNMLLLEESN GYKIFGKTGWAMDI
OXA24 DNFWLVGPLKITPVQEVNFADDLAHNRLPFKLETQEEVKKMLLIKEVN GSKIYAKSGWGMGV
OXA45 THSWLGSSLKISPEGQVRFVRDLLSAKLPASKDAQQMTVSILPHFAAG DWAVQGKTGTGSFIDAR
OXA46 GDYWIDGNLKISAHEQILFLRKLYRNQLPFKVEHQRLVKDLMITEAGR SWILRAKTGWEG--
OXA51 DNFWLVGPLKITPQQEAQFAYKLANKTLPFSPKVQDEVQSMLFIEEKN GNKIYAKSGWGWDV
OXA57 DRAWIGSSLQISPLEQLEFLGKMLDRKLPVSPTAVDMTERIVESTTLADGTVVHGKTGVSYPLLAD
OXA58 DKFWLKGPLTITPIQEVKFVYDLAQGQLPFKPEVQQQVKEMLYVERRG ENRLYAKSGWGMAV
OXA85 DKFWLEGPLKISAMEQVKLLNLLSQSKLPFKLENQEQVKDITILEKKD DFILHGKTGWATDN
OXA143 DNFWLVGPLKITPIQEVNFADDFANNRLPFKLETQEEVKKMLLIKEFN GSKIYAKSGWGMDV
OXA427 TQAWLSSSLAISPQEQARFLGKLVSGKLPVSAQTLQHTANILRQPDID GWQIHGKTGTGYPKLLD
AFD1 THSWLGASLTISPVEQVGFLRRLLGGNLPFSRDAQAKTRAIMPVFDAPESWAVHGKTGTGYMRDEK
ATD1 THSWLGASLTVSPVEQVGFIRRLLAGNLPVSRDAQAKTRAIVPVFDAPESWSVHGKTGTGFMRDEK
CPD1 DTFWLDNSLQISPDEELGFVKKLYFDQLPFHKVTMQNVRQVMLMEKKP EYELSYKTGMGFSG
LoxA9 THAWLSSSLSISPTEQIQFLQKIIYKKLPVSQKAYTMTKNIMYIQELPGGWKLYGKTGTGRQLTKD
NOD1 DRFWLEGPLQITPKQQIEFLQRLHRKELPFSQRTLDLVQDIMIYERTP NYVLRGKTGWAASV
STD1 NSFWIDRSLKISPEEQIDFLINLYEEKFMLSEKTYKIVKDIMINEKTP EYTLRGKTGWGREG
Gram-positive
BAT2 DQFWLQSSLKISPLEEKDFIEHLYKEDLPFDKPIMKTVKRMMIQEEGD HYTLYGKTGTRLT-
BSU2 DQFWLQSSLTISPLEQETFLEKLAKEELPFDKPVMKIVKRMMIQEEGD HYTLYGKTGTRLT-
BPU1 DQFWLSSTLRISPIEQVRFLKQLYEETLPFDLKNMRTVKRMMVQEEEK HATLYGKTGSG---
α8’
CDD1 GDLKNGKGHLNGFWLESSLQISPKEQVQTMAKIFEGDTNFKKEHINILRDIMAIDVNDANINVYGKTGTGFDEK
CDD10 GDLKNGKGHLNGFWLESSLQISPKEQVQTMAKIFEGDTNFKKEHINILRDIMKIDVNDKNINVYGKTGTGFDEK
consensus FWL KTG
-153a -153b -153c -153d -153e -153f -153g -153h -153i -153j -201a -215a -215b -215c
DBL number 160 170 180 190 200 210
type-III
β10 β11 α11
Gram-negative
OXA48 EPKIGWWVGWVELDD NVWFFAMNMDMPTS DGLGLRQAITKEVLKQEKIIP 265
OXA1 RTLQNGWFEGFIISKSGHKYVFVSALTGNLGS NLTSSIKAKKNAITILNTLNL 276
OXA2 --RMGWWVGWVEWPT GSVFFALNIDTPNRM DDLFKREAIVRAILRSIEALPPNPAVNSDAAR 275
OXA10 SNPGVAWWVGWVEKET EVYFFAFNMDIDNE SKLPLRKSIPTKIMESEGIIGG 266
OXA23 KPQVGWLTGWVEQPDGKIVAFALNMEMRSEM -PASIRNELLMKSLKQLNII 273
OXA24 TPQVGWLTGWVEQANGKKIPFSLNLEMKEGM -SGSIRNEITYKSLENLGII 275
OXA45 GAKAPLGWFIGWATHEE RRVVFARMTAGGKKG EQPAGPAARDAFLKALPDLAKRF 264
OXA46 --RFGWWVGWIEWPT GPVFFALNIDTPNRT DDLFKREAIARAILRSIDALPPN 266
OXA51 DPQVGWLTGWVVQPQGNIVAFSLNLEMKKGI -PSSVRKEITYKSLEQLGIL 274
OXA57 GTRDWARGSGWFVGWIVRGK QTLVFARLTQDERKQ PVSAGIRTREAFLRDLPRLLAAR 269
OXA58 DPQVGWYVGFVEKADGQVVAFALNMQMKAGD -DIALRKQLSLDVLDKLGVFHYL 280
OXA85 IVVPIGWFVGWIETSD NIYSFAINLDISDS KFLPKREEIVREYFKNINVIK 261
OXA143 TPQVGWLTGWVEKSNGEKVAFSLNIEMKQGM -PGSIRNEITYKSLENLGII 275
OXA427 GSLDRDQQIGWFVGWASKQD QKLIFVHTVIQKPGKQ FASLRAREEVFAALPEQLKKL 264
AFD1 GNPDRNRPFGWFVGWAEREG QHIVFARLRVSDKPS NEPLGPAVRDAFLRDIPRLAVHR 274
ATD1 GNPDRSRPFGWFVGWAEREG QHIVFARLRVADKPS SEPLGPAVRDAFLRDIARLAVHR 273
CPD1 PKTIGWITGWIEENG HPSFFVLNIETENKS LDMRTVRMNILRNLLTDAGYFKGMK 267
LoxA9 KSQKLPLQHGWFVGWIEKDE RVITFAKHIADSKEN TTFASFRAKNDTLIQLFNLINELEK 266
NOD1 TPNIGWFVGYLEQNN NVYFFATNIDIRNN DDAAARIEVTRRSLKALGLL 283
STD1 AENIIWYVGYIEAKE NVYFFAVRIINASEERNSYLLDFRKQLTMMAFRELGIIN 264
Gram-positive
BAT2 DYGLGWFVGFITTDN HSYIFVTNVDA--- -SGTAAKNITTDILKKYHLITN 267
BSU2 DMGLGWFVGFIKTEH GSYVFVTNVDD--- -SGTKAKNITVDILKKYGLITS 267
BPU1 -SDIGWYVGFIKHEH KTYILATNIKG--- -TGIEAKDITYRILKKYHLMEASV 262
CDD1 NKRVDAWFVGMLEREG DTYYFAIKSDDSNKE ITGPKVKEIAINIIKKYYS 309
CDD10 NKRVDAWFVGMLEREG DTYYFAIKSDDSNKE ITGPKVKEIAINIIKKYYSVREGAAL 294
consensus GWxxGW
-215d -215e -215f -215g -215h -230a -244a -244b -244c -244d
DBL number 220 230 240 250 260 270

**Figure S2. Structure-based sequence alignment of 25 class D β-lactamases.** This alignment was used to generate a standard DBL residue numbering and secondary structure annotation scheme. Amino acids in helices are colored blue and those in β-strands are colored red, and the secondary structure annotation is given at the top of each block. The P-loop and the Ω-loop are highlighted in gray boxes, and the locations of the two different types (II and III) are indicated. The new DBL numbering scheme is shown at the bottom of each block. Insertions in sequences relative to OXA-48 are indicated by gaps in the alignment, and the number of the last residue before the insertion with an alphabetical suffix is given at the bottom of each block. Deletions in sequences relative to OXA-48 are indicated by dashes. The seven conserved class D sequence motifs noted in Figure 1 are highlighted in light green, and their consensus sequences are indicated at the bottom. Residues in the motifs which are >95% conserved across all class D β-lactamases are given in bold in the consensus sequence. Amino acids colored gray at the N-termini of some sequences are parts of the mature enzymes which were not modeled in the structures which were used for superposition against OXA-48 prior to manual adjustment of the alignment.

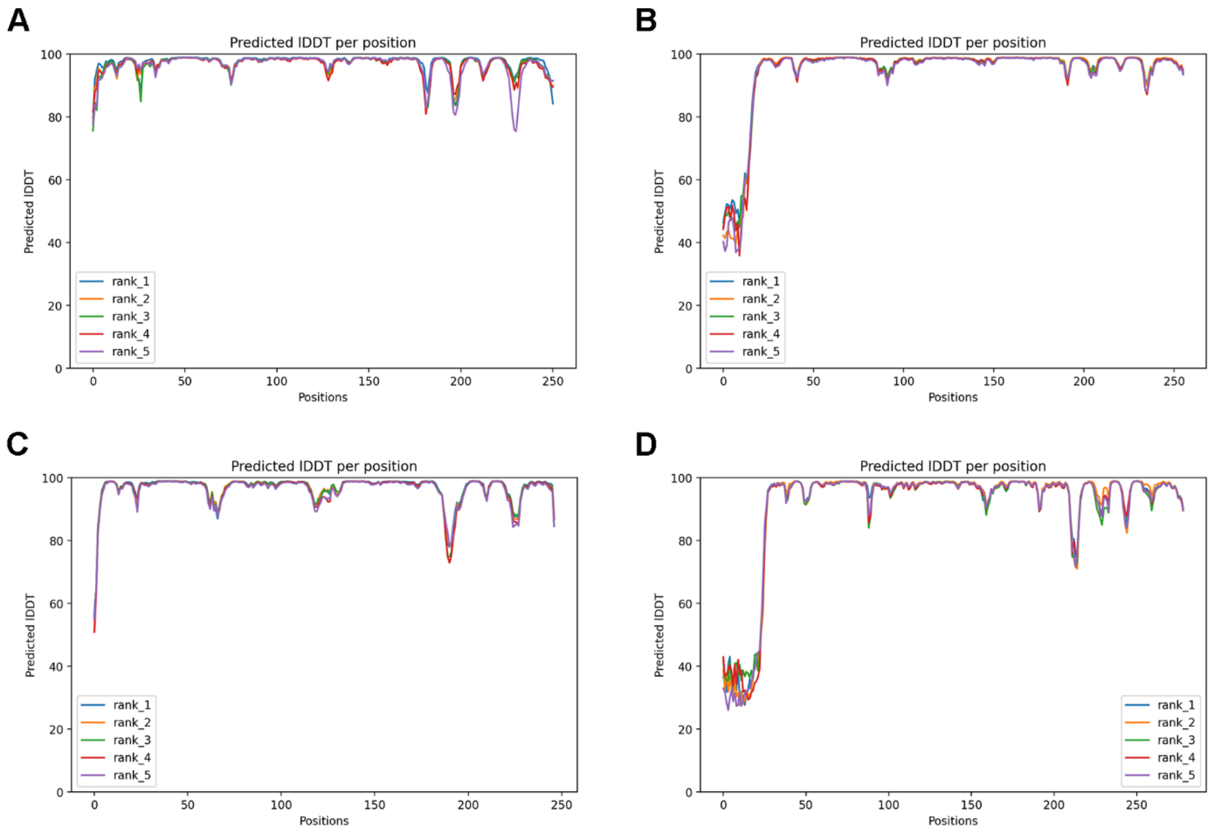


**Figure S3. Plots of predicted LDDT for test targets.** The pLDDT is given per residue position as a function of residue number for the five ranked AlphaFold2 models for the four class D test targets. **(A)** OXA-1024. **(B)** OXA-213. **(C)** OXA-919. **(D)** ClosD1. The first ~25 residues of the predicted ClosD1 structure had substantially lower pLDDT scores (<40) for all five ranked models and these residues were removed prior to superposition against OXA-48.

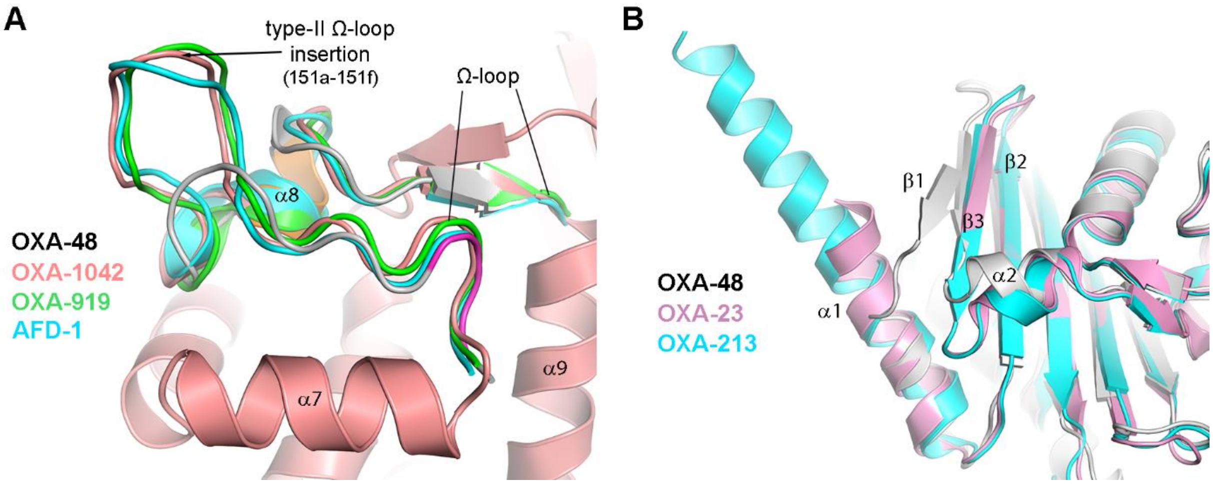


**Figure S4. DBL test targets (A)** Superposition of the type-II Ω-loop of OXA-919 (green) and AFD-1 (cyan), and the type-I loop in OXA-48 (gray), onto the predicted OXA-1042 model (salmon). The DBL numbering for the loop extension is indicated. **(B)** Superposition of the predicted OXA-213 model (cyan) and OXA-23 (pink) onto OXA-48 (gray), showing the extended helix α1 and the lack of a strand β1 in the first two structures.

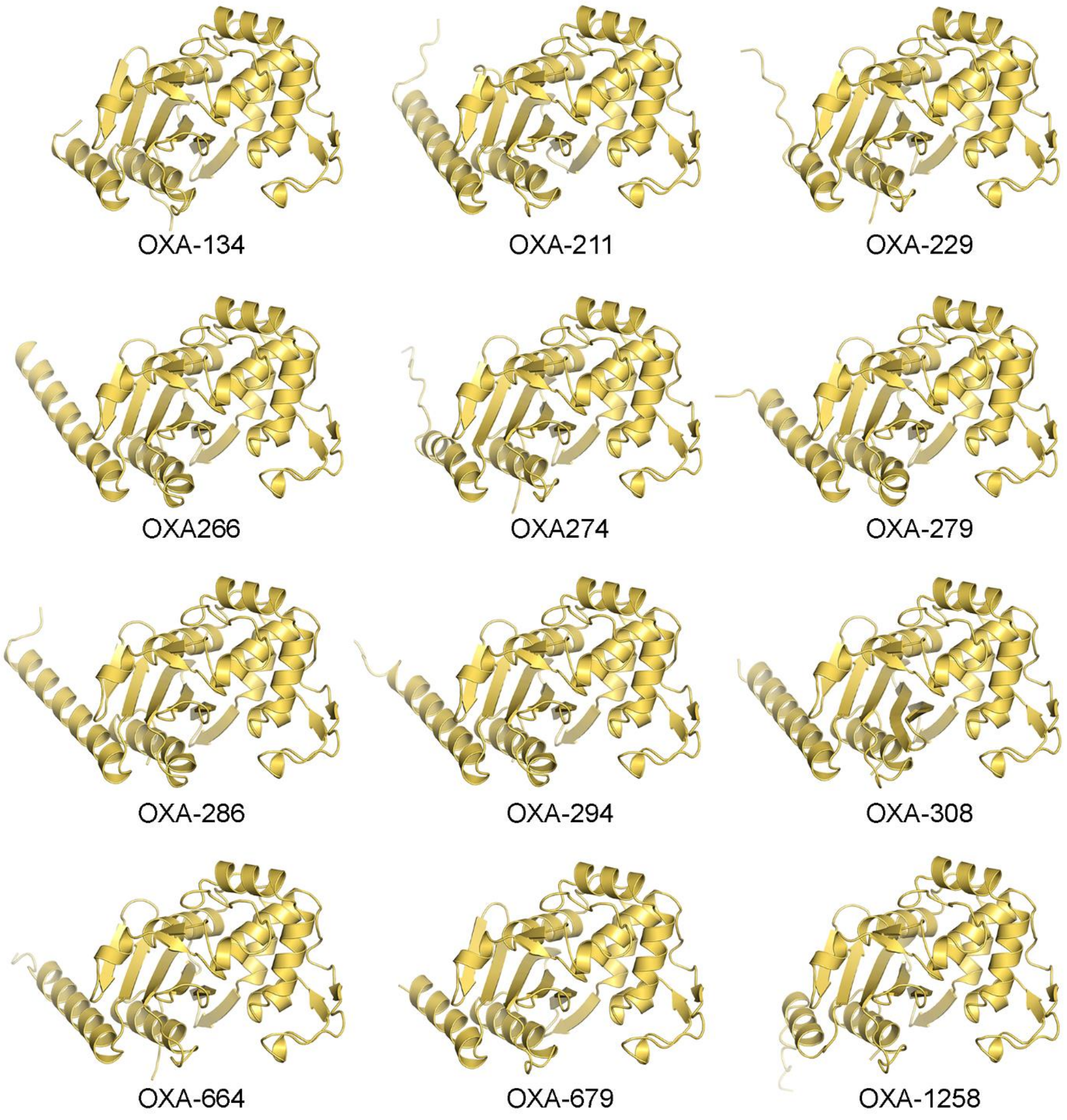


**Figure S5.** AlphaFold2[1] predicted structures of the eponymous sequences of twelve *Acinetobacter* families where there is currently no structural information (excluding OXA-213 which was used as a test target in this paper). The signal sequences of all enzymes were identified with DeepSig[2] and removed prior to AlphaFold2 prediction. The pLDDT scores for the predicted structures were, in order from top left; 95.4, 95.8, 94.8, 96.9, 95.1, 96.2, 96.4, 95.9, 95.4, 92.8, 95.1 and 91.9.

**Table S1. The 50 OXA enzyme families**[a]

| > 10 members | 5-10 members | < 5 members |
|---|---|---|
| OXA-1 (13) | OXA-22 (7) | OXA-5 (2) |
| OXA-2 (32) | OXA-46 (6) | OXA-9 (2) |
| OXA-10 (64) | OXA-55 (9) | OXA-20 (2) |
| OXA-12 (11)[a] | OXA-58 (9) | OXA-42 (4)[b] |
| OXA-23 (54) | OXA-60 (7) | OXA-62 (3) |
| OXA-24 (15) | OXA-214 (6) | OXA-198 (4) |
| OXA-48 (68) | OXA-243 (10) | OXA-258 (3) |
| OXA-50 (104) | OXA-274 (6) | OXA-266 (2) |
| OXA-51 (390) | OXA-548 (6) | OXA-279 (3) |
| OXA-61 (49) | OXA-679 (7) | OXA-308 (3) |
| OXA-63 (23) | OXA-1258 (6) | OXA-372 (3) |
| OXA-114 (23) | | OXA-427 (2) |
| OXA-134 (31) | | OXA-464 (4) |
| OXA-143 (12) | | OXA-493 (2) |
| OXA-184 (39) | | OXA-664 (3) |
| OXA-211 (18) | | OXA-732 (2) |
| OXA-213 (106)[c] | | OXA-1043 (2) |
| OXA-229 (26) | | OXA-1149 (4) |
| OXA-286 (17) | | |
| OXA-294 (13) | | |
| OXA-504 (12) | | |

[a] A family is determined to be a grouping of two or more sequences with greater than 90% identity and they are named according to the first enzyme identified. The number in each family are given in parentheses. The families colored red have at least one member whose structure has been determined.
[b] Represented by the OXA-57 structure (99% identity with OXA-42)
[c] Includes the OXA-270 sub-family (85-90% identity to OXA-213)

**Table S2. Class D β-lactamase secondary structure assignment from literature reports**[a]

| OXA-1 | OXA-10 | OXA-10 | OXA-13 | OXA-23 | OXA-24 | OXA-46 | OXA-48 | OXA-51 | OXA-57 | OXA-58 | OXA-143 | OXA-145 | OXA-163 | OXA-163 | OXA-181 | OXA-245 | OXA-935 | BPU-1 | CDD-1 | BSU-2 BAT-2 | consensus |
|---|---|---|---|---|---|---|---|---|---|---|---|---|---|---|---|---|---|---|---|---|---|
| 3 | 4 | 5 | 6 | 7 | 8 | 9 | 10 | 11 | 12 | 13 | 14 | 15 | 16 | 17 | 17 | 17 | 18 | 19 | 20 | 21 | |
| β1 | β1 | β1 | β1 | | | β1 | β1 | | | | | β1 | β1 | β1 | β1 | β1 | β1 | | β0 | | β1 |
| α1 | α1 | α1 | α1 | α1 | α1 | α1 | α1 | α1 | | α1 | α1 | | α1 | α1 | α1 | α1 | α1 | α1 | α1 | α1 | α1 |
| β2 | β2 | β2 | β2 | β1 | β1 | β2 | β2 | β1 | β1 | β1 | β1 | β2 | β2 | β2 | β2 | β2 | β2 | β1 | β1 | β1 | β2 |
| β3 | β3 | β3 | β3 | β2 | β2 | β3 | β3 | β2 | β2 | β2 | β2 | β3 | β3 | β3 | β3 | β3 | β3 | β2 | β2 | β2 | β3 |
| α2 | α2 | α2 | | α2 | α2 | α2 | α2 | α2 | | α2 | α2 | α1 | α2 | α2 | α2 | α2 | | α2 | α2 | α2 | α2 |
| β4 | βa1 | β4 | | β3 | | | | β3 | | β3 | β3 | | | β4 | β4 | β4 | | β3 | β3 | β3 | β4 |
| α3 | α3 | α3 | α3 | α3 | α3 | α3 | α3 | α3 | α1 | α3 | α3 | α2 | α3 | α3 | α3 | α3 | α3 | α3 | α3 | α3 | α3 |
| β5 | βb1 | | | | | | | β4 | | β' | β4 | | | | | | | β4 | β4 | β4 | β5 |
| | | | | | | | $3_{10}$ | $3_{10}$ | α2 | $3_{10}$ | $3_{10}$ | α3 | α4 | | | | | $3_{10}$ | $3_{10}$ | $3_{10}$ | α4 |
| β6 | βb2 | | | | | | | β5 | | β'' | β5 | | | | | | | β5 | β5 | β5 | β6 |
| α4 | α4 | α4 | α4 | α4 | α4 | α4 | α4 | α4 | α3 | α4 | α4 | α4 | α5 | α4 | α4 | α4 | α4 | α4 | α4 | α4 | α5 |
| α5 | α5 | α5 | α5 | α5 | α5 | α5 | α5 | α5 | α4 | α5 | α5 | α5 | α6 | α5 | α5 | α5 | α5 | α5 | α5 | α5 | α6 |
| α6 | α6 | α6 | α6 | α6 | α6 | α6 | α6 | α6 | | α6 | α6 | α6 | α7 | α6 | α6 | α6 | α6 | α6 | α6 | α6 | α7 |
| α7 | α7 | | | | | | α7 | α7 | α5 | α7 | α7 | | α8 | α7 | α7 | α7 | | | α7 | α7 | α8 |
| β7 | βa2 | β5 | | β4 | | | | β6 | | β4 | β6 | | | β5 | β5 | β5 | | β6 | β6' | β6' | β7 |
| α8 | α8 | α7 | α7 | α7 | α7 | α7 | α8 | α8 | α6 | α8 | α8 | α7 | α9 | α8 | α8 | α8 | α8 | α7 | α8 | α8 | α9 |
| α9 | α9 | α8 | α8 | α8 | α8 | α8 | α9 | α9 | α7 | α9 | α9 | α8 | α10 | α9 | α9 | α9 | α9 | α8 | α9 | α9 | α10 |
| β8 | β4 | β6 | β6 | β5 | β3 | β4 | β4 | β7 | β3 | β5 | β7 | β4 | β4 | β6 | β6 | β6 | β4 | β7 | β7 | β7 | β8 |
| β9 | β5 | β7 | β7 | β6 | β4 | β5 | β5 | β8 | β4 | β6 | β8 | β5 | β5 | β7 | β7 | β7 | β5 | β8 | β8 | β8 | β9 |
| β10 | β6 | β8 | β8 | β7 | β5 | β6 | β6 | β9 | β5 | β7 | β9 | β6 | β6 | β8 | β8 | β8 | β6 | β9 | β9 | β9 | β10 |
| β11 | β7 | β9 | β9 | β8 | β6 | β7 | β7 | β10 | β6 | β8 | β10 | β7 | β7 | β9 | β9 | β9 | β7 | β10 | β10 | β10 | β11 |
| α10 | α10 | α9 | α9 | α9 | α9 | α9 | α10 | α10 | α8 | α10 | α10 | α9 | α11 | α10 | α10 | α10 | α10 | α9 | α10 | α10 | α11 |

[a] References are given for each structure in the second row.

**Table S3. Insertions and deletions in class D structures relative to OXA-48**

| | Number of residues | DBL position | Location |
|---|---|---|---|
| Insertions | 1 | 28 | β1-α1 loop |
| | ≤ 3 | 50 | β2-β3 loop |
| | 1 | 54 | β3[a] |
| | 2 | 57 | β3-α2 loop[b] |
| | 1 | 91 | α3-β5 loop |
| | ≤ 2 | 101 | P-loop |
| | ≤ 6 | 151 | Ω-loop (type-II) |
| | 10 | 153 | Ω-loop (type-III)[c] |
| | 1 | 201 | β8-β9 loop |
| | ≤ 8 | 215 | β9-β10 loop[d] |
| | 1 | 230 | β10-β11 loop |
| | ≤ 4 | 244 | β11-α11 loop[d] |
| Deletions | 1 | 29 | β1-α1 |
| | 2 | 37-38 | α1-β2 loop |
| | 1 | 50, 51 | β2-β3 loop |
| | 2 | 60-61 | β3-β4 loop[e] |
| | 1 | 87 | α3-β5 loopf |
| | 3 | 213-215 | β9-β10 loop |
| | 4 | 242-245 | β11-α11 loop |

[a] This insertion is present in OXAs -1, -45, -57, -427, AFD-1, ATD-1 and LoxA9.
[b] This insertion is present only in *Acinetobacter* OXAs.
[c] Present in the enzymes from *C. difficile*.
[d] These loops have a variable length in several enzymes.
[e] Helix α2 is missing in enzymes with this deletion.
[f] This deletion is present in OXA-1 and the *Acinetobacter* OXAs.

**Table S4. DBL scheme test targets.**

| Category[a] | Enzyme | Host | Signal peptide[b] | pLDDT / pTM | *rmsd* (Å)[c] |
|---|---|---|---|---|---|
| 1 | OXA-1042 (OXA-1)[d] | *A. veronii* | 1-25 | 97.4 / 0.944 | 1.6 |
| 2 | OXA-213 | *A. baumannii* | 1-17 | 95.1 / 0.904 | 1.3 |
| 3 | OXA-919 | *B. pseudogrignonensis* | 1-28 | 96.6 / 0.938 | 1.8 |
| 4 | ClosD1[e] | *Clostridium* sp. CH2 | 1-26 | 91.3 / 0.877 | 1.7 |

[a] Category 1 = sequences from an OXA family where a structure is known; Category 2 = sequences from an OXA family with no known structural representative; Category 3 = other Gram-negative enzyme sequences or OXA orphans; Category 4 = Gram-positive enzyme sequence.

[b] The signal peptide was predicted using the DeepSig server[2] and was removed from the sequence prior to the pairwise sequence alignment and AlphaFold2 calculation.

[c] The *rmsd*s were calculated relative to OXA-48 (PDB code 3HBR).

[d] OXA-1042 was chosen because it is the newest annotated members of the OXA-1 family.

[e] The *Clostridium* sequence was downloaded from the Bacterial and Viral Bioinformatics Resource Center (BV-BRC).
